# Identification of ethanol analgesia quantitative trait loci and candidate genes in BXD recombinant inbred mouse lines

**DOI:** 10.1101/2024.06.17.599372

**Authors:** Walker D. Rogers, Alyssa White, M. Imad Damaj, Michael F. Miles

**Author notes:** **Corresponding authors:** Michael Miles, MD, PhD, and M. Imad Damaj, PhD **E-mail:**.

## Abstract

Alcohol consumption produces acute analgesic effects, and people experiencing pain conditions may drink alcohol to alleviate discomfort. However, tolerance to the analgesic properties of alcohol could prompt escalating consumption and dependence. Both nociception and alcohol-induced analgesia are under significant genetic control. Understanding the genetic architecture of these processes could inform better treatment options for people with pain conditions. This study aims to identify quantitative trait loci (QTL) driving variation in ethanol-induced analgesia across BXD recombinant inbred mouse lines. Male and female mice from 62 BXD strains received ethanol or saline oral gavage for five days and were tested for hot plate (HP) latency at baseline, Day 1, and Day 5. QTL mapping of HP phenotypes identified a significant provisional QTL on chromosome 17 for Day 1 HP latency in mice receiving ethanol. An additional highly suggestive QTL was present on chromosome 9 for the difference in pre- and post-ethanol thermal nociception. Candidate genes within QTL support intervals were provisionally identified using HP phenotypic correlations to transcriptomic database, expression QTL analysis, and other bioinformatics inquiries. The combined behavioral and bioinformatic analyses yielded strong ethanol analgesia candidate genes, specifically *Myo6*. Thus, the results of this genetic study of ethanol-induced analgesia in BXD mouse strains may contribute significantly to our understanding of the molecular basis for individual variation in the analgesic response to acute ethanol.

## Introduction

Alcohol use disorder (AUD) and pain conditions are significant health concerns that affect millions of people in the United States each year. Both are associated with increased risk of mortality, with alcohol-related deaths exceeding 170,000 per year in the United States, and both AUD and pain represent a burden of hundreds of billions of dollars annually in healthcare costs and productivity loss in the United States[1–3]. These are related public health problems, as people with pain conditions are significantly more likely to endorse AUD than the general public[4]. Alcohol confers analgesic effects in a dose-dependent fashion in humans in a laboratory setting, and people with pain conditions often self-report drinking alcohol to manage pain symptoms[5, 6]. However, excessive alcohol consumption is associated with AUD, cardiovascular disease, cirrhosis, type 2 diabetes, and many cancers, among other aspects of alcohol consumption detrimental to public health[7].

The relationship between alcohol use and pain is complex, and the bidirectional pathway between these two health conditions often leads to co-occurring alcohol consumption and pain conditions. Excessive alcohol consumption can lead to painful health outcomes. For example, an estimated 44% of chronic alcohol users develop peripheral neuropathy, which can be characterized by paresthesia, numbness, and pain[8]. Alcohol misuse is also a primary cause of both acute and chronic pancreatitis, a painful inflammation of the pancreas[9]. Prolonged alcohol consumption causes withdrawal-induced hyperalgesia, in which pain sensitivity is significantly increased relative to basal pain thresholds, in both humans and rodent models [10, 11]. Repeated cycles of pain, intoxication, and hyperalgesia may contribute to increasing alcohol consumption and the development of AUD[12]. Complicating our understanding of pain- and alcohol-related phenotypes is the fact that these phenomena are complex and often under some degree of genetic control. Peripheral neuropathy can result from either environmental or hereditary causes, for example, and there is considerable interpersonal variation in alcohol analgesia due to genetic and environmental factors[13–15]. Elucidating the molecular genetic basis for ethanol analgesia is critical for understanding both alcohol use disorder and pain conditions.

Recently, studies in rodent model organisms have described specific roles for opioid receptors and GABAergic signaling pathways in alcohol analgesia[16–18]. However, these systems do not account for all the variation in alcohol-induced analgesia. To identify quantitative trait loci (QTL) modulating alcohol analgesia on a genome-wide scale, we performed thermal nociception assays on the BXD panel of genetically informative recombinant inbred mice derived from the C57BL/6J and DBA/2J parental strains. This panel is valuable for characterizing the genetic component of complex behaviors, and it has been used extensively for addiction research for over 20 years. BXD lines have previously been used to characterize QTL underlying ethanol consumption, anxiolytic-like ethanol responses, ethanol sensitivity, and morphine-induced analgesia[19–21].

We assessed male and female mice from 62 BXD strains for hot plate latency at baseline, post-acute ethanol, and post-repeat ethanol timepoints. Our studies identified multiple significant and suggestive post-ethanol QTL. Genes located in the support intervals of these QTL were investigated using complementary bioinformatic and systems genetics approaches that leveraged historical brain region-specific expression correlation data, phenomic overlap with other ethanol or analgesia traits, and the presence of cis-acting expression QTL. Our results yield a novel list of provisional candidate genes for the regulation of post-ethanol thermal nociception.

## Materials and Methods

### Animals

Male (n = 175) and female (n = 171) BXD mice from 62 strains were obtained from Jackson Laboratory (Bar Harbor, ME) at 8 weeks of age. Mice were housed 4-5 per cage and had access to rodent chow and water. All animals were acclimatized to vivarium housing for one week prior to experimentation and underwent gavage habituation with water for 3 days prior to ethanol studies. Mice were ordered with randomization of strain and sex so as to avoid batch effect confounds. To avoid sex-specific confounding of nociception assays, no male laboratory staff were present for behavioral testing.

### Nociception assay

Hot plate testing protocols were adapted from White et al. (2020), which showed 1) 2.0 g/kg oral gavage ethanol produces maximum antinociception at 30 minutes post-administration 2) this dose of ethanol shows no locomotor effects in B6 mice and a small locomotor effect in D2 mice 3) baseline hot plate testing (as in the current design) does not interfere with post-gavage thermal nociception (i.e. post-saline Day 1 hot plate latency should be indistinguishable from Baseline)[18]. After acclimation the testing room overnight, mice were tested for baseline 55°C hot plate latency (Thermojust Apparatus, Columbus, OH) (Figure S1). The latency until the first nocifensive behavior (licking paw, shaking paw) was recorded, and a cutoff time of 20 seconds was established to minimize risk of tissue damage. Two hours after baseline latency, mice received oral gavage of 2.0 g/kg ethanol (v/v) in water (Sigma-Aldrich, St. Louis, MO) or volumetric equivalent saline. Thirty minutes post-gavage, mice were again assayed for latency on a 55°C hot plate. To assay tolerance to ethanol analgesia, mice received daily 0.5 g/kg ethanol or saline oral gavage for the following three days and were not behaviorally assayed during this time. On the final day of the experiment, mice again received 2.0 g/kg ethanol or saline and were assayed for hot plate latency 30 minutes post-gavage. The basal, post-acute ethanol, and post-repeated ethanol timepoints are respectively referred to as Baseline, Day 1, and Day 5. The difference between Baseline and Day 1 is referred to as Difference.

### QTL mapping

Quantitative trait locus mapping was performed with the R package qtl2[22]. QTL mapping results were validated using Hayley Knott regression on GeneNetwork (https://genenetwork.org). To identify possible sex-specific QTL, interval mapping was conducted on strain means for each treatment group and timepoint for male mice, female mice, and collapsed across sex. The laboratory technician conducting the nociception assay was used as a covariate to control for interpersonal variation. Kinship matrices were constructed using the “leave one chromosome out” method to account for relatedness among mice. BXD strain genotypes covering 7320 markers across the genome were downloaded from https://gn1.genenetwork.org/dbdoc/BXDGeno.html. Significance thresholds for LOD scores were calculated with 1000 permutations of the phenotypic and genotypic data, and support intervals for bQTL were derived from a 1.5-LOD drop around the peak marker of the corresponding QTL. Genome scans were performed on raw data or square root-transformed data if transformation produced a normal distribution.

### Statistical analyses

A three-way ANOVA was performed to measure the effects of sex, genotype, and ethanol treatment on hot plate latency at each timepoint. Sex was not a significant predictor of hot plate latency at any measurement, and subsequent analyses were collapsed across sex. A two-way ANOVA examining the effects of timepoint and ethanol treatment group on hot plate latency was performed, and group means were compared using Tukey’s test of honestly significant differences. All calculations were performed using R, and the R scripts used for analysis and data visualization are publicly available (https://osf.io/9ph3g/). Heritability estimates and their standard errors for behavioral phenotypes were calculated with a linear mixed model using the est_herit() function in the R package qtl2, which used a kinship matrix to account for relatedness between BXD strains.

### Systems genetics

Rather than only using QTL positional information to identify provisional candidate genes for behavioral phenotypes, we utilized existing behavioral and transcriptomic datasets on BXD mice that are contained within the GeneNetwork resource. We uploaded the hot plate latency data and and queried genes with basal expression profiles correlated with post-ethanol hot plate latency (pre-frontal cortex, GN135; nucleus accumbens, GN156[23]; and midbrain, GN381[24]). The same gene expression microarray datasets were queried for genes with significant expression quantitative trait loci (eQTL) within the support interval around the bQTL. Our search defined an eQTL as a marker within 2Mb of the target gene with a minimum likelihood ratio statistic (LRS) of 20. For eQTL queries, results with polymorphisms in the probe set were excluded to prevent false positives when probe sequence information was available. Hot plate phenotypes were also correlated with existing BXD traits uploaded to GeneNetwork. The top 10000 trait correlations were filtered for descriptions including the terms “hot plate,” “analgesia,” “nociception,” “ethanol,” and “pain.”

## Results

### Oral ethanol administration acutely decreases thermal nociception

A total of 356 mice underwent hot plate latency testing at Baseline, Day 1 acute post-ethanol, and Day 5 post-repeated ethanol timepoints. A three-way ANOVA of sex, timepoint, and treatment on hot plate latency revealed main effects for timepoint and treatment but no main effect of sex or any significant interaction terms for sex (Table 1). Male and female results were thus collapsed for each treatment-timepoint group.

**Table 1:** ANOVA of hotplate behavioral data.

| Term | DF | SS | Mean Sq | F | Pr(>F) |
| --- | --- | --- | --- | --- | --- |
| Timepoint | 2 | 304.7 | 152.4 | 11.9 | 7.815e-06* |
| Sex | 1 | 6.52 | 6.52 | 0.509 | 0.4757 |
| Treatment | 1 | 63.55 | 63.55 | 4.961 | 0.02614* |
| Timepoint:Sex | 2 | 1.129 | 0.5647 | 0.04409 | 0.9569 |
| Timepoint:<br>Treatment | 2 | 269 | 134.5 | 10.5 | 3.057e-05* |
| Sex:Treatment | 1 | 0.1368 | 0.1368 | 0.01068 | 0.9177 |
| Timepoint:<br>Sex:Treatment | 2 | 36.31 | 18.16 | 1.417 | 0.2428 |
| Residuals | 1026 | 13142 | 12.81 | NA | NA |

ANOVA results showing the significant effects of timepoint and ethanol treatment on hot plate latency. Sex was not a significant predictor of thermal nociception for any timepoint or ethanol treatment group. Significant (<0.05) raw p-values are denoted with asterisks.

Acute ethanol administration significantly prolonged hot plate latency in mice, although there was no difference between the treatment groups at Baseline or Day 5 timepoints (Figure 1A). Interestingly, mice in the vehicle group exhibited reduced thermal nociception on Day 5 compared to the previous two timepoints (Day 5 – Baseline p_adj_: 3.6e-4; Day 5 – Day 1 p_adj_: 6.3e-4). This finding is consistent with other studies that have examined repeated hot plate testing in rodent models[25, 26]. The decreased sensitivity may be a result of learned behavioral responses or stress-induced analgesia. For ethanol-receiving mice, average hot plate latency increased significantly from 8.52 (S.E. +/- 0.244) s to 10.3 (S.E. +/- 0.317) s from Baseline to Day 1 measurements. Day 1 and Day 5 latencies for ethanol-receiving mice were not significantly different from each other (p_adj_ = 0.245) and Day 5 responses did not differ in control versus ethanol-receiving animals (p_adj_ = 0.717).

**Figure 1.**
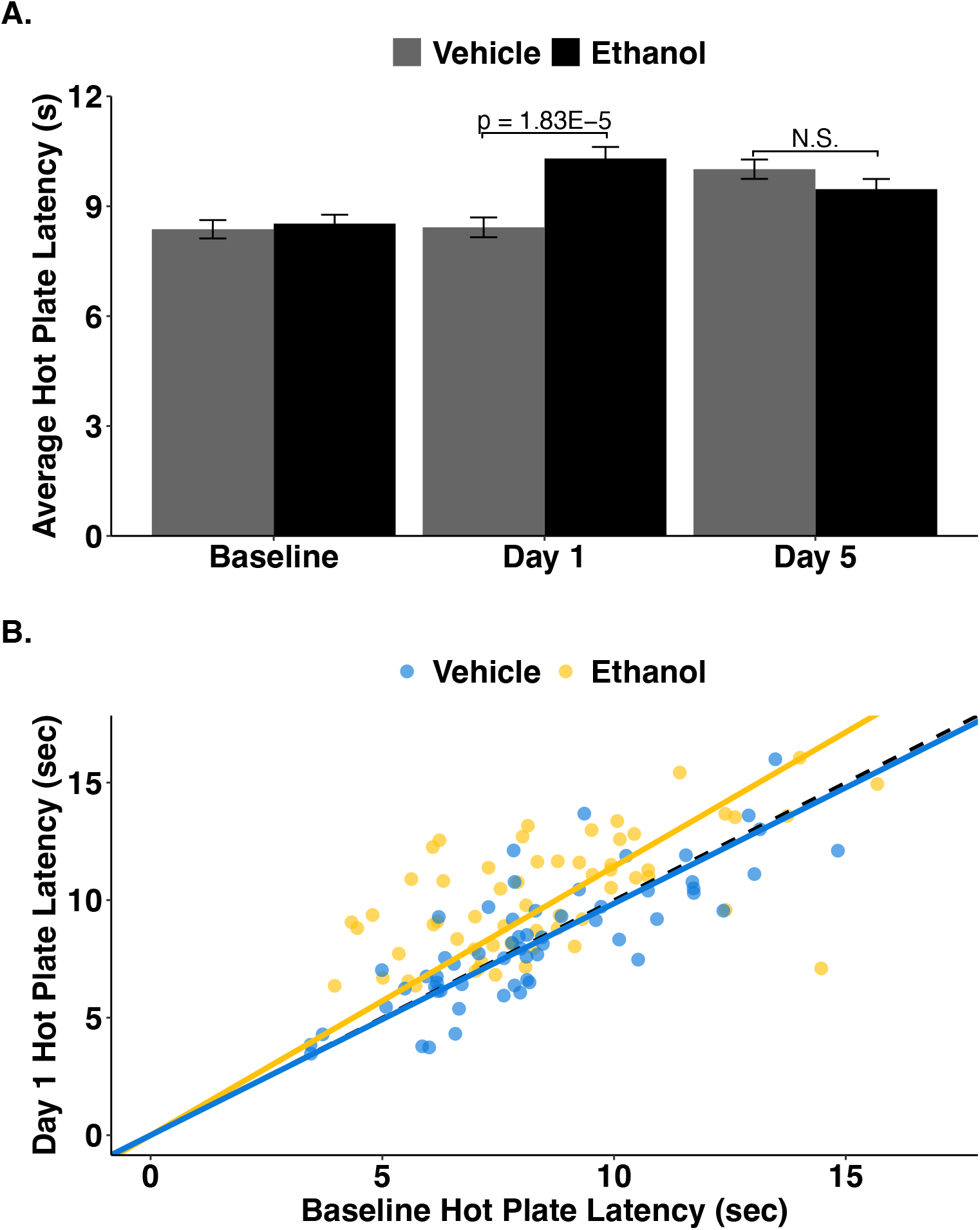
Mean hot plate latencies in BXD mice. A) Mean hot plate latency (+/- 1 SE) in seconds at each timepoint for vehicle- and ethanol-receiving mice. Acute ethanol significantly increases hot plate latency in BXD mice. B) A scatterplot depicts BXD strain mean Baseline and Day 1 hot plate latencies in seconds on the x and y-axes, respectively. Regression lines for each treatment were set to an intercept at the origin, and the dashed diagonal (slope = 1) represents a hypothetical perfect correlation between Baseline and Day 1 latency.

The Baseline and Day 1 hot plate latencies for mice receiving saline showed a highly significant positive correlation, r(164) = .69, p = 2.2e-16. This correlation was attenuated in the ethanol group, presumably due to inter-strain variation in ethanol analgesia, r(167) = .55, p = 7.2e-15. A scatterplot visualizing Baseline and Day 1 hot plate latencies for strain means in each treatment group shows vehicle-receiving mice clustered tightly around the diagonal, indicating little variation from initial testing results (Figure 1B). Ethanol-receiving mice are clustered above the diagonal, reflecting the significant decrease in thermal nociception caused by acute ethanol across the population, except for several strains showing possible increased thermal nociception following ethanol.

### Post-ethanol hot plate latency is not confounded by locomotor activity

Basal differences in locomotor activity (LMA) between BXD strains has the potential to influence results from thermal nociception assays. Because the hot plate assay is based on the latency of a mouse to move in response to thermal stimuli, mice that exhibit higher basal LMA may be incorrectly judged to have lower hot plate latency. To prevent spurious antinociception results, the LMA of each mouse was measured in photocell activity cages following Day 1 and Day 5 hot plate assays. To characterize the relationship between post-ethanol LMA and hot plate latency, the Pearson correlation between these variables was calculated (Figure 2). Post-acute ethanol (Day 1) hot plate latency was not significantly correlated with either Day 1 initial locomotor activity (r(167) = -.08, p = .27) or Day 1 total locomotor activity (r(167) = -.09, p = .27) as measured by the photocell activity cage. The difference between Baseline and Day 1 latency in ethanol-receiving mice was not correlated with Day 1 initial locomotor activity (r(167) = −0.13, p = 0.09), although it was modestly negatively correlated with Day 1 total locomotor activity (r(167) = −0.17, p = 0.03). However, a one-way ANOVA of Day 1 total LMA on the Difference phenotype yields only a small effect size (η^2^ = 0.03) of this LMA phenotype on ethanol group Day 1 – Baseline Difference. In mice receiving saline gavage, the Day 1 – Baseline Difference is similarly correlated with Day 1 Total LMA, although the locomotor activity again has a small effect size on the nociception phenotype (η^2^ = 0.03).

**Figure 2.**
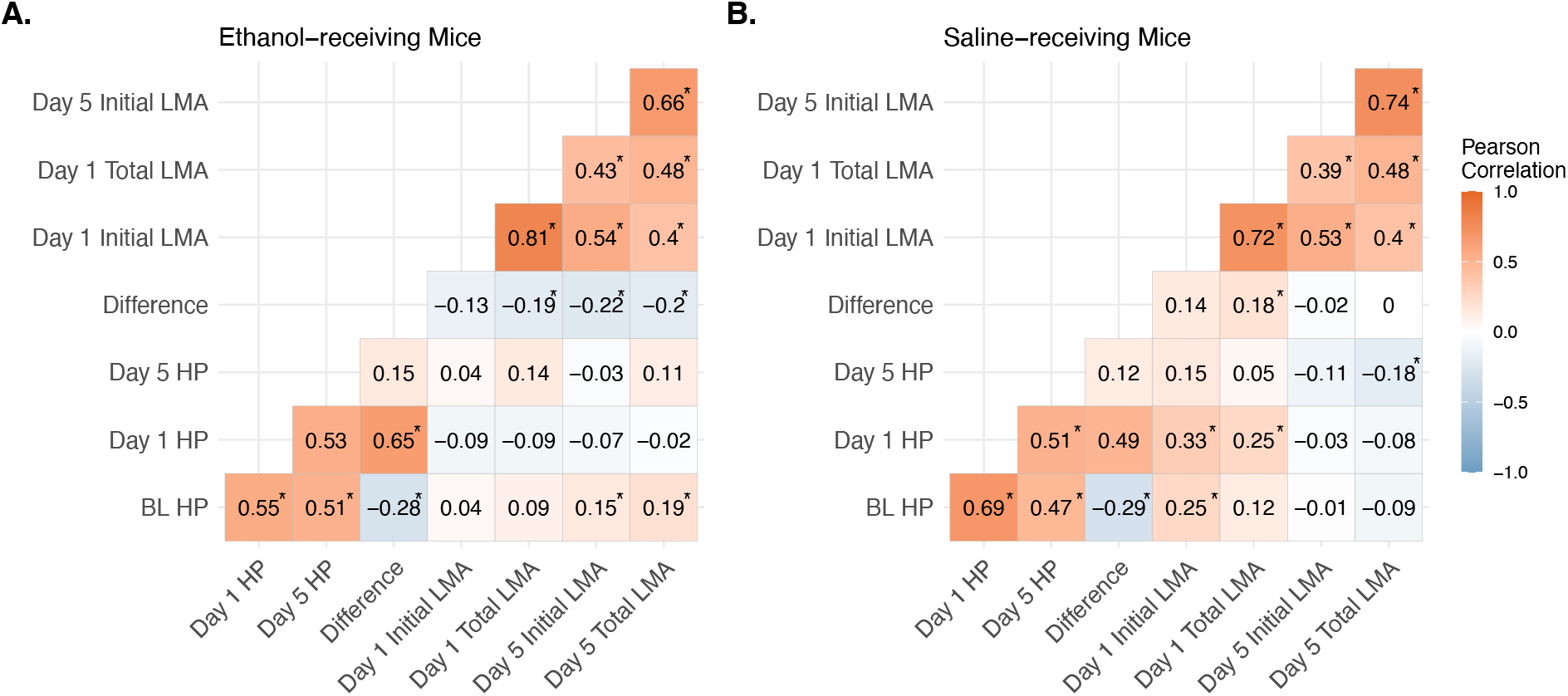
Correlograms for locomotor activity and hot plate latency phenotypes in ethanol-receiving mice (2A) and saline-receiving mice (2B). Asterisks denote a significant uncorrected p-value < 0.05 for the correlation.

### QTL interval mapping yields novel loci regulating thermal antinociceptive properties of ethanol

For each timepoint and treatment group, we performed interval mapping in the R package qtl2 using a linear mixed model to account for relatedness among BXD strains. Mapping was performed for individual sexes and also collapsed across sex to identify both sex-specific QTL and leverage greater statistical power from pooled samples (Table S1). Sex-specific analysis did not identify any significant QTLs, but multiple highly suggestive or suggestive QTLs were identified within either sex for latencies at various timepoints, treatments and analysis of Day1-Baseline latencies for ethanol treatment. However, due to the ANOVA results above showing no significant sex or sex*treatment differences, we chose to focus on analysis of data collapsed across sexes for increased statistical power.

We identified a significant QTL (Pehlq1; **p**ost-**e**thanol **h**ot plate **l**atency QTL 1) on chromosome 17 for Day 1 post-acute ethanol hot plate latency (peak LOD = 4.17, p<0.05) with a 1.5-LOD support interval of 3.67Mb (Chr17: 64.40-68.07Mb) (Figure 3A). A query of this interval on Mouse Genome Informatics database yields 23 protein-coding genes (Table S2). Mice with B6 alleles at the peak marker had reduced hot plate latency after acute ethanol than mice with D2 alleles at this locus (Welch two sample t-test, p<0.0001) (Figure 3B). To identify differences in the QTL landscape between treatment groups for this phenotype, we performed the same mapping for mice that received saline gavage. At the Day 1 timepoint, there was no QTL on chromosome 17 for hot plate latency in vehicle-receiving mice, and the maximum LOD score at this locus was less than 0.5 (p>>0.05) (Figure S2). These results suggest an ethanol-specific linkage between thermal nociception responses and genetic variants in the Chr 17 7.5Mb interval.

**Figure 3.**
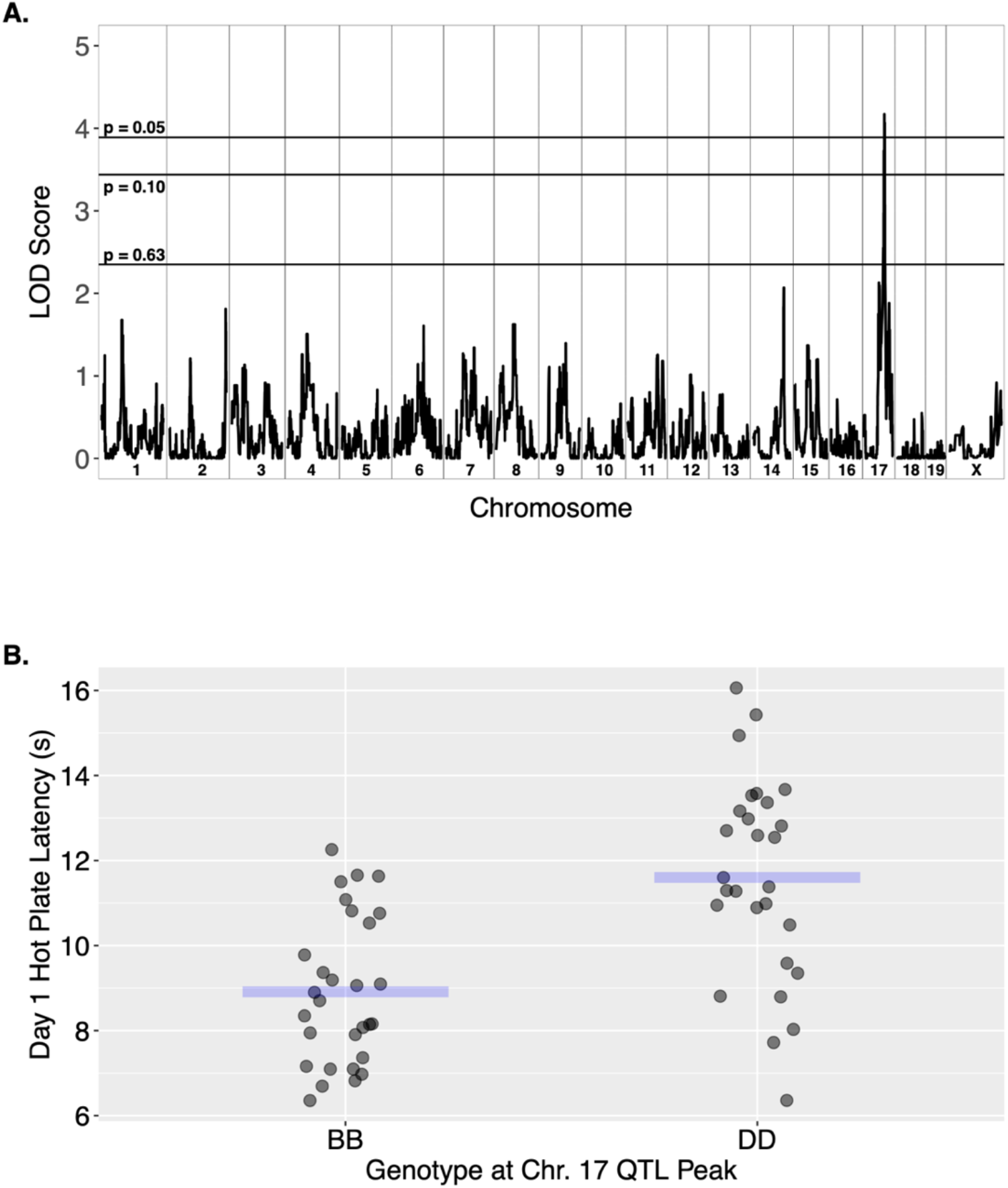
QTL analysis for Day 1 post-ethanol hot plate latency. A) The genome scan of square root-transformed Day 1 post-ethanol hot plate latency yields a significant QTL peak with a LOD score >4 on chromosome 17. B) Mice with DBA2/J alleles at this locus exhibit increased post-ethanol latency compared with mice with C57BL6/J alleles (Welch two sample t-test, p<0.0001).

For a complementary approach to assessing ethanol analgesia, we also investigated the difference between Day 1 and basal hot plate latency (Difference). The mean Difference in the 171 vehicle-receiving mice was 0.05 seconds, and the mean Difference in 175 ethanol-receiving mice was 1.78 seconds. The Difference phenotype is useful, as it is robust to basal differences in thermal nociception among BXD strains. Using this phenotype as input for interval mapping, we identified a highly suggestive QTL on chromosome 9 (Etadq1; **Et**hanol **a**nalgesia **d**ifference QTL 1) with a peak LOD score of 3.096 (p<0.10) and a 1.5-LOD interval of 73.5-83.3 Mb (Figure 4). This interval contains 57 protein-coding genes (Table S2).

**Figure 4.**
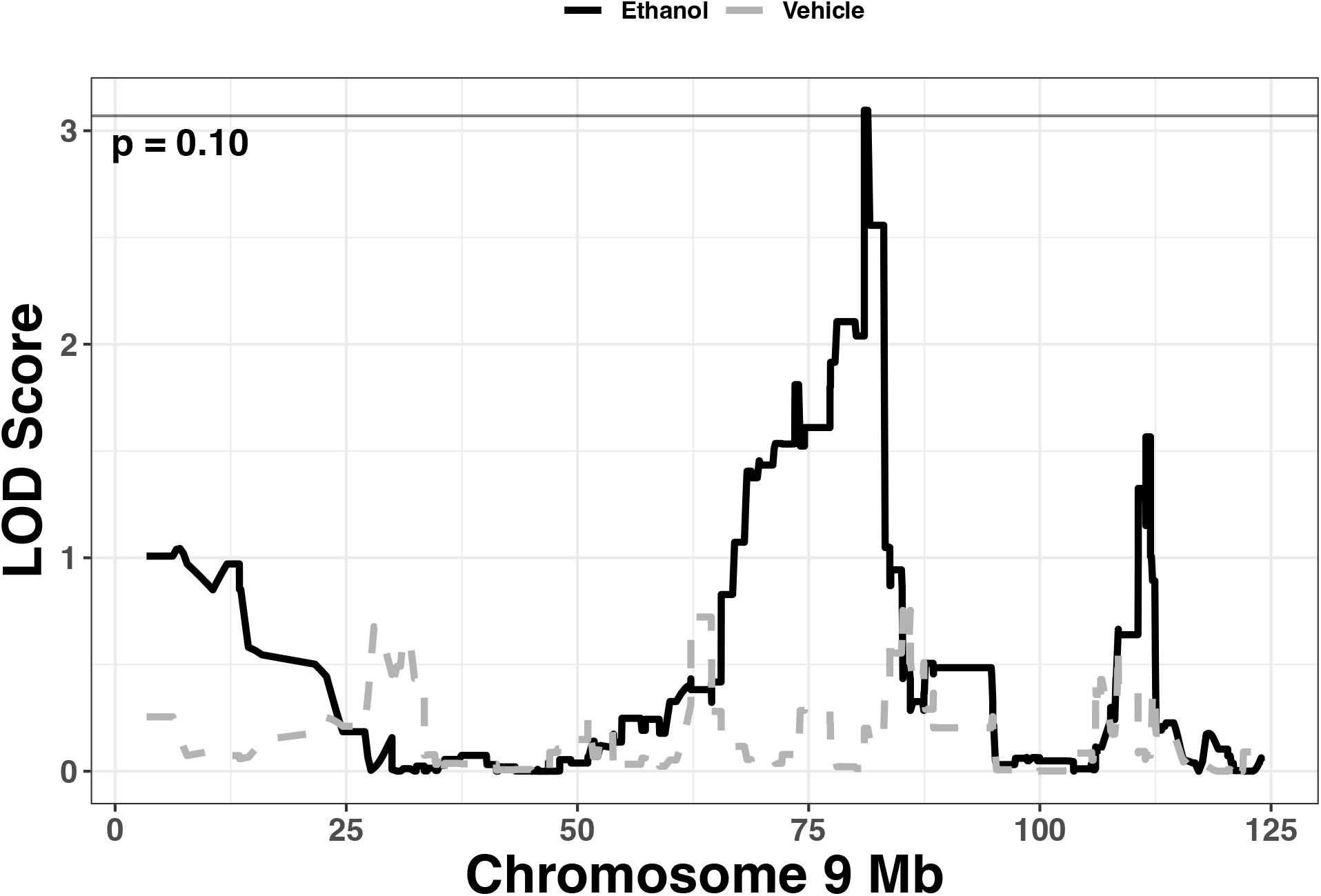
Chromosome 9 QTL for difference between pre- and post-treatment hot plate latency. A highly suggestive ethanol-specific QTL (p < 0.10) on chromosome 9 was identified for the difference in pre- and post-treatment thermal nociception. No QTL was observed at this locus when performing a genome scan for the Difference phenotype in saline-receiving mice (dashed grey line).

### Brain region-specific expression QTL overlap with genes in ethanol analgesia behavioral QTL

In the human GWAS literature, almost 90% of tagged SNPs are located in non-coding intronic or intergenic regions of the genome, implying that these variants are driving differential gene expression to modulate complex traits[27]. To identify genetic differences in gene expression, we characterized expression quantitative trait loci (eQTL) for genes in the support interval of ethanol analgesia QTL. We queried GeneNetwork expression databases for basal gene expression in prefrontal cortex, nucleus accumbens, and midbrain to identify genes in the chromosome 17 *Pehlq1* QTL and the chromosome 9 *Etadq1* support intervals with significant cis-acting eQTL (Table S3). The chromosome 17 bQTL support interval contained nine genes with significant cis-eQTL, one of which were represented in more than one brain region (*Tmem232*). The chromosome 9 bQTL support interval included 29 genes with putative cis-eQTL, with *Elovl5* exhibiting cis-eQTL in all three regions. These genes all had cis-acting variants driving variable expression levels between the BXD progenitor strains and are therefore strong candidates for ethanol analgesia candidate genes. Filtering for presumptive false positive eQTL due to polymorphisms in microarray probe sets yielded 7 genes across the two QTL support intervals with significant cis-acting eQTL (Table 2).

**Table 2:**
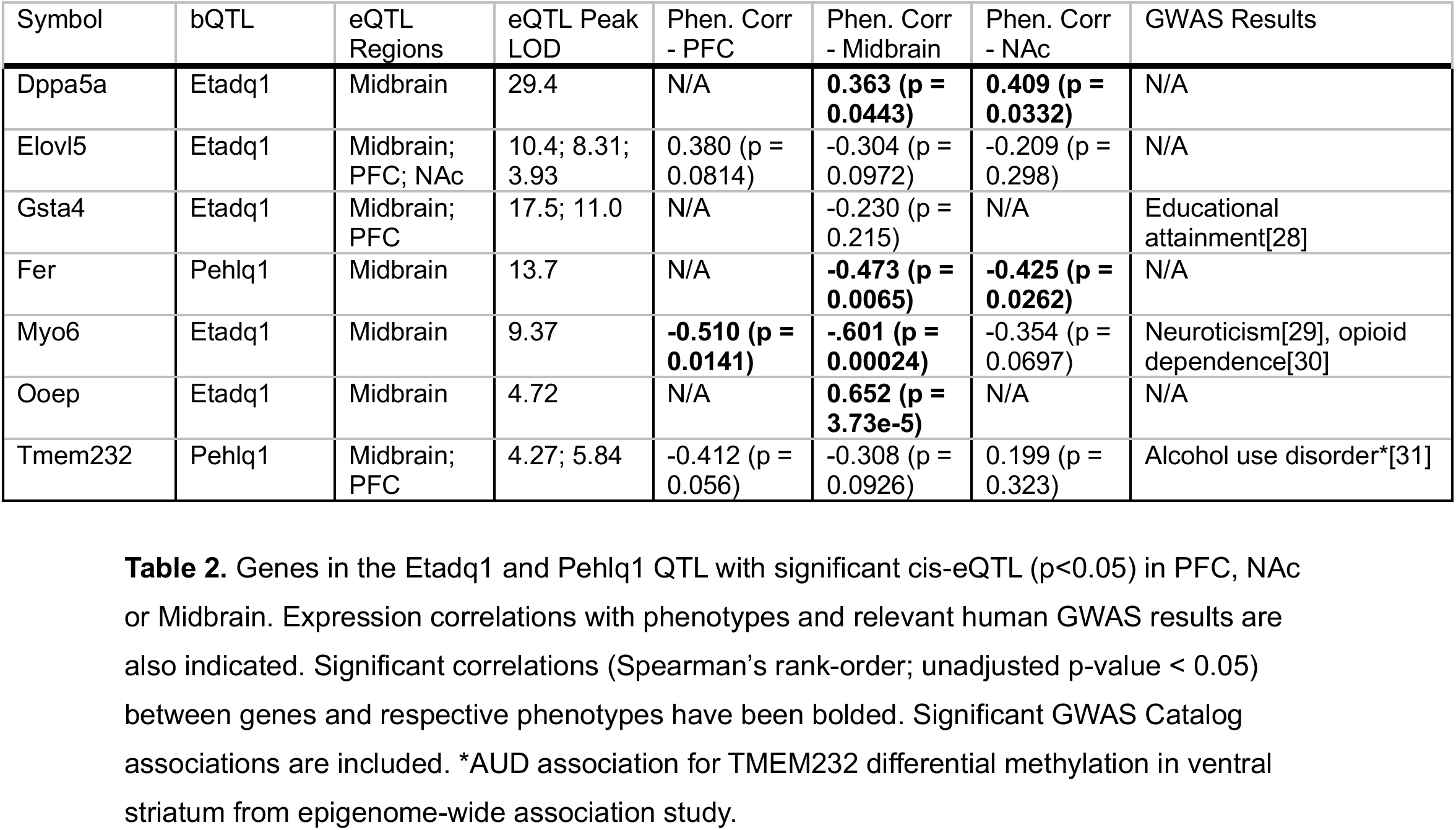
Candidate genes in QTL support intervals.

### Brain regional BXD gene expression patterns correlate with hot plate phenotypes

To identify genes that may be regulating ethanol analgesia responses in mice, post-ethanol hot plate latency and Day 1 – Baseline difference traits were correlated with basal brain gene expression data from BXD mice in prefrontal cortex, nucleus accumbens, and midbrain in GeneNetwork (Table 2; GN traits #135, 156, 381)[23, 24, 32]. Significant correlations may indicate causative roles for the genes in the target phenotype, so the top 10,000 phenotype-expression correlations from each brain region were downloaded for analysis. All brain gene expression-hot plate phenotype correlates are available in Table S4. For genes in the *Pehlq1* interval, our query yielded 23 expression correlations across the three brain regions that were significant at an uncorrected p-value of 0.05. The correlation of *Tmem232* and *Fer* expression with post-ethanol hot plate latency was significant in more than one brain region. For genes in the *Etadq1* support interval, Oocyte expressed protein (*Ooep*) in the midbrain yielded the strongest correlation of any gene-phenotype pair (p = 3.73e-5), although its expression was not significantly correlated in other brain regions. Midbrain-specific *Myo6* expression was significantly negatively correlated (p = 6.3e-4) with the difference in Baseline and post-ethanol Day 1 hot plate latency (Figure S2). The chromosome 9 QTL support interval contained 11 genes with significant uncorrected Spearman rank correlations. Table 2 shows that *Dppa5a*, *Fer*, *Myo6* and *Ooep* all have significant eQTLs and expression correlation results in the same expression data.

### Hot plate phenotypes are correlated with historic BXD ethanol and nociception traits

The latency phenotypes for each timepoint and the Day 1 – Baseline measurement were also analyzed by genetic correlation with publicly available BXD databases on GeneNetwork to identify related biological phenotypes. The full list of phenotype-phenotype correlations can be viewed in Table S5. Results were filtered by searching the top 10,000 Spearman rank correlations for the phrases “ethanol,” “alcohol,” “nociception,” “analgesia,” and “locomotor activity.” The difference between Day 1 and Baseline thermal nociception in control group mice was strongly correlated with the hot plate latency of saline-receiving animals (r(16) = −0.64, p = 0.0065) from another nociception study (Record ID 10020)[33]. A highly suggestive chromosome 12 QTL for basal hot plate latency in male mice (Table S1) overlapped the confidence interval for a previously reported thermal nociception QTL (Record ID 10897)[34] located in the same region on chromosome 12. Additionally, the baseline hot plate latency of ethanol-receiving mice in our study was significantly correlated (Figure S3) with the basal hot plate latency from a 1993 study by Mogil et al[35].

In ethanol-receiving mice, the Day 1-Baseline Difference was also significantly correlated (rho(42) = −0.38, p = 0.01) with basal thermal nociception of female mice from a high throughput screening of phenotypes in the BXD panel by Philip et al. (Record ID 11640)[36].

Although the mice in the comparison study did not receive ethanol, the observed correlation is informative. Strains exhibiting a high difference between latency measurements tended to have lower basal hot plate latencies in the comparison study, which may be due to mice with higher Baseline latencies reaching the 20s hot plate cutoff time after acute ethanol. The Day 1 – Baseline difference phenotype in the ethanol group was also significantly correlated with measures of voluntary ethanol consumption under the drinking in the dark paradigm (IDs 13576, 13578, 20312 and 20314)[37]. Interestingly, the difference in latency for the ethanol group was nominally correlated with serotonin levels in the blood of female mice (ID 17970)[38]. The gene closest to the peak marker of *Etadq1* is *Htr1b*, a serotonin receptor implicated in numerous psychiatric disorders, including alcohol and opioid use disorders, suicidal behavior, and major depressive disorder[39, 40].

### Alcohol, neuropsychiatric phenotypes in human GWAS findings from syntenic bQTL intervals

We queried the JAX Synteny Browser for the support intervals of the *Pehlq1* and *Etadq1* QTLs to identify possible consilience with related human traits. These genomic regions were entered into the human GWAS catalog to identify variants with significant associations with alcohol or pain traits from the literature. All associations with p-values and mapped genes from these queries can be found in Table S6. The chromosome 9 *Etadq1* interval is syntenic with regions on human chromosomes 6 (53.00-55.88, 73.36-85.68Mb) and 15 (51.68-54.68Mb). Within this region are numerous variants significantly associated with alcohol phenotypes and psychiatric traits. Eleven variants with genome-wide significant associations with drinks per week were mapped to SUMO specific peptidase 6 (SENP6). Variants associated with drinks per week, schizophrenia, smoking initiation, and bipolar disorder were mapped to synaptosome associated protein 91 (SNAP91). The chromosome 17 *Pehlq1* interval maps to regions on human chromosomes 5 (109.5-110.7Mb) and 18 (2.57-8.83; 9.10-9.96Mb), and it features variants associated with opioid analgesic sensitivity (DLGAP1)[41], schizophrenia (MAN2A1)[42], chronic pain (TMEM200C)[43], and post-traumatic stress disorder symptomatology (DLGAP1)[44].

### Ethanol metabolism correlates

To characterize the possible role of ethanol pharmacokinetics in our phenotypes, we correlated Day 1 and Difference phenotypes from the ethanol group with historic blood ethanol concentration (BEC) datasets in GeneNetwork. We revealed mixed results with previous studies. For Day 1 latencies, some studies yielded significant uncorrected correlations with blood ethanol concentrations (IDs 11710, 11967, 10347)[36], while there was no correlation with other BEC measurement data (IDs 19470, 10177)[20]. The Difference phenotype yielded similar results when correlated with BEC data. In one comparison, the Day 1 – Baseline difference is significantly correlated (p = 0.02) with ethanol sensitivity as measured by BEC at loss of balance during a dowel test (ID 10347)[45]. However, other BEC measurement reports yielded no correlation with the Difference data (ID 11710, 10178)[20]. The highest coefficient of determination for any of the analgesia phenotype-BEC correlations was 0.21, indicating that most of the variation in hot plate latency is attributable to factors other than ethanol metabolism.

## Discussion

Problematic alcohol use and pain disorders are overlapping health problems that significantly impact the quality of life of millions of people each year. Because of the prevalence of alcohol self-administration in patients with pain, it is critical to understand the genetic and molecular mechanisms underlying ethanol analgesia. In this study we conducted the first genome-wide QTL mapping study to identify genetic loci linked with variation in ethanol-induced analgesia in acute thermal pain test, using BXD mice as a model system. Our study yielded a significant QTL for post-ethanol hot plate latency on mouse chromosome 17 and a highly suggestive QTL for ethanol analgesia response on mouse chromosome 9. By characterizing genes located within these QTL, we have curated a list of candidate genes correlated with ethanol analgesia and related behavioral phenotypes. Evidence for possible causal roles in the antinociceptive properties of ethanol were derived from brain gene expression-hot plate latency correlations, human GWAS literature, behavioral QTL mapping, and the presence of *cis*-eQTL. Together this work has confirmed the important genetic influence on ethanol’s analgesic properties and identified promising candidate genes for future study.

Our behavioral examination of basal and post-ethanol hot plate latency revealed a significant antinociceptive effect of ethanol, congruent with rodent[16, 18] and human[5] analgesia literature. There was no difference in hot plate latency between ethanol and control groups by Day 5 nociception testing, though, due to an increase in latency in vehicle-receiving mice. Other studies have reported similar results after repeated thermal nociception assays in rodents[25], and future experiments to characterize tolerance to ethanol analgesia will need to account for this acclimatization.

This study reinforced the heritability of complex phenotypes for nociception, ethanol responses, and analgesia. Estimates of heritability for Baseline and Day 1 hot plate latency in both saline controls and ethanol-receiving mice were significant (i.e., heritability estimate +/- standard error did not overlap 0) (Table S7). The heritability estimate for the Day 1 – Baseline Difference measurement was not significant in the ethanol group of mice, but this may be a function of the number of phenotypes used to calculate this input introducing increased variance.

As noted in Results, we focused our analysis efforts on data derived from pooling sexes due to the ANOVA results showing no sex or sex x treatment differences. However, Table S1 does identify multiple sex-specific suggestive or highly suggestive QTLs that may warrant future study. In particular, a broad region on chromosome 12 contained suggestive or highly-suggestive QTLs across basal, Day 1 or Day 5 timepoints in either sex and in pooled data.

When analyzing data pooled across sexes, behavioral QTL mapping identified a region on mouse chromosome 17 significantly linked with Day 1 hot plate latency in ethanol-receiving mice. This QTL likely represents an interaction of genetic influences on basal hot plate latency and ethanol-induced analgesia, although no suggestive or significant QTLs were found in this region for analysis of basal responses (Table S1; see below). As the first timepoint after ethanol administration, this measurement provided useful information regarding thermal nociception in the context of acute ethanol analgesia. With the large number of strains assayed, as well as the inclusion of advanced recombinant inbred BXD strains, the support interval for this QTL was less than 4 Mb, thus aiding candidate gene prioritization. This significant Day 1 latency QTL appeared in ethanol-receiving mice collapsed across sex, but it is also a suggestive QTL in the male ethanol-receiving group. Although sex was not a significant predictor of any hot plate latency or difference phenotype, the differing results in male and female mice may be caused by quantitative or qualitative genetic differences or increased environmental variance in female animals. Importantly, these QTL are derived from acute thermal nociception measures, and more research will be needed to address the genetic architecture of chronic pain exposure.

The difference between baseline hot plate latency and Day 1 post-ethanol hot plate latency was calculated as an additional measurement of ethanol analgesia. As this phenotype reflects the change from baseline to post-ethanol latency, it should be robust to differences in basal thermal sensitivity across BXD strains. This chromosome 9 QTL peak (*Etadq1*) was highly suggestive in pooled data and suggestive in female ethanol-receiving mice but not male mice, again underscoring the importance of sex differences in the genetic influences on ethanol analgesia. Significant QTLs for Day 5 – baseline hot plate latency in ethanol-receiving mice were identified (Table S1), but interpretation and validation of these loci will require further studies given the increase in hot plate latency observed in vehicle groups at that time point.

To further leverage the large BXD dataset on ethanol analgesia generated in this study, historic BXD datasets were queried to identify significant correlations between our nociception variables and other phenotypes relevant to nociception, analgesia, and ethanol or other drugs. The Day 1 post-ethanol hot plate latency phenotype was significantly correlated with other hot plate latency results, including baseline thermal nociception before a forced-swim test (GN ID 10418)[35]. Although the comparison study did not administer ethanol to the mice, it is unsurprising to see strong concordance with our post-ethanol latencies since we also found a significant correlation between Day 1 post-ethanol latency and basal hot plate latencies in this study (Figure 2). These results reflect upon the relatively large role of strain-specific basal thermal sensitivity versus ethanol analgesia per se and suggest that mechanisms of ethanol analgesia impinge directly upon those mediating basal nociception, as might be expected.

We coupled the above bioinformatic analyses with a search of human GWAS results of positional candidate genes from the *Pehlq1* and *Etadq1* bQTL. Human genome regions syntenic to the *Etadq1* interval returned variants associated with neuroticism and opioid use disorder (*MYO6*), schizophrenia (*SNAP91*), calcium levels (*CD109*), and alcohol consumption (*SENP6*, *SNAP91*). The genetic overlap between alcohol use disorder and other psychiatric traits, including schizophrenia and neuroticism[46], is well-characterized, and it’s possible that the variants underlying these traits have a pleiotropic effect on ethanol analgesia responses.

Calcium channel activity is known to regulate morphine analgesia[47], and CD109-specific modulation of calcium levels may contribute to differential ethanol analgesia across the BXD lines in our study.

Myosin VI (*MYO6*) is a gene encoding an actin-dependent motor protein that is broadly expressed in the brain[48], and it has classically been associated with hearing loss and deafness due to its role in the structure of hair cells in the inner ear[49]. More recently, *Myo6* has been implicated in synaptic plasticity through its role in the endocytosis of AMPA receptors in Purkinje cells[50, 51], and a variant in *MYO6* was nominally correlated (p = 5.24e-7) with opioid use disorder in individuals of European ancestry[30]. In the field of alcohol research, *Myo6* has been shown to be downregulated in myelin-forming oligodendrocyte cells in mouse cerebellum following chronic-plus-binge ethanol exposure [52]. A synonymous single nucleotide variant in *Myo6* was one of only 12 loci in the rat genome significantly associated with ethanol consumption in the alcohol-preferring (P) rodent model and the high-alcohol drinking (HAD) replicated lines[53]. Given the recent findings in model organism and human alcohol literature, the genomic position in the novel *Etadq1* ethanol analgesia QTL, the correlation between midbrain *Myo6* expression and ethanol-mediated antinociception, and the significant *cis*-eQTL regulating expression in PFC, we hypothesize a role for *Myo6* in regulating ethanol analgesia in mice.

Excessive alcohol use is a significant cause of mortality worldwide, with the World Health Organization estimating that over 5% of deaths annually are attributable to alcohol-related causes[54]. Alcohol is inexpensive, accessible, and analgesic, and many people with pain conditions report alcohol use to treat pain[6, 55]. As alcohol analgesia is a heritable trait, identifying the molecular genetic mechanisms responsible for antinociception is critical to preventing the misuse of alcohol as a painkiller. In this first reported mouse genetic analysis of ethanol analgesia, we identified novel QTL linked with post-ethanol thermal nociception (*Pehlq1*) and ethanol antinociceptive responses (*Etadq1*). With a complementary systems genetics approach, we offer supporting evidence of *Myo6* as an ethanol analgesia candidate gene. Further mechanistic studies will be needed to establish a causal role for this gene in the analgesic action of acute ethanol, as well as for extending analysis of the additional suggestive QTLs and candidate genes identified in this report.

## Acknowledgments

The authors would like to thank Emily Legge for her contributions to the hot plate assays and animal care.

## Data Availability Statement

The data and R scripts used for analysis and generation of figures are available at https://osf.io/9ph3g/.

## Abbreviations

bQTL: behavioral quantitative trait loci
B6: C57BL/6J mice
BXD: C57BL/6J X DBA/2J recombinant inbred mice
Chr: chromosome
eQTL: expression quantitative trait loci
D2: DBA/2J mice
FDR: false discovery rate
HP: hot plate
LOD: logarithm of odds
Mb: megabase
NAc: nucleus accumbens
PFC: prefrontal cortex
QTL: quantitative trait loci
SI: support interval
SNP: single nucleotide polymorphism

---

**Figure S1.**
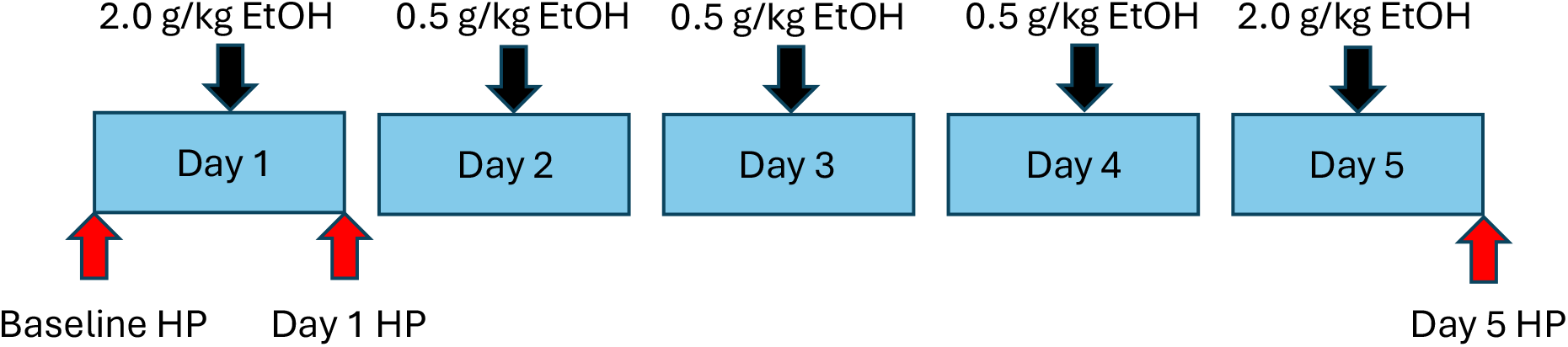
Timeline of ethanol analgesia behavioral testing protocol. EtOH = ethanol; HP = hot plate.

**Figure S2.**
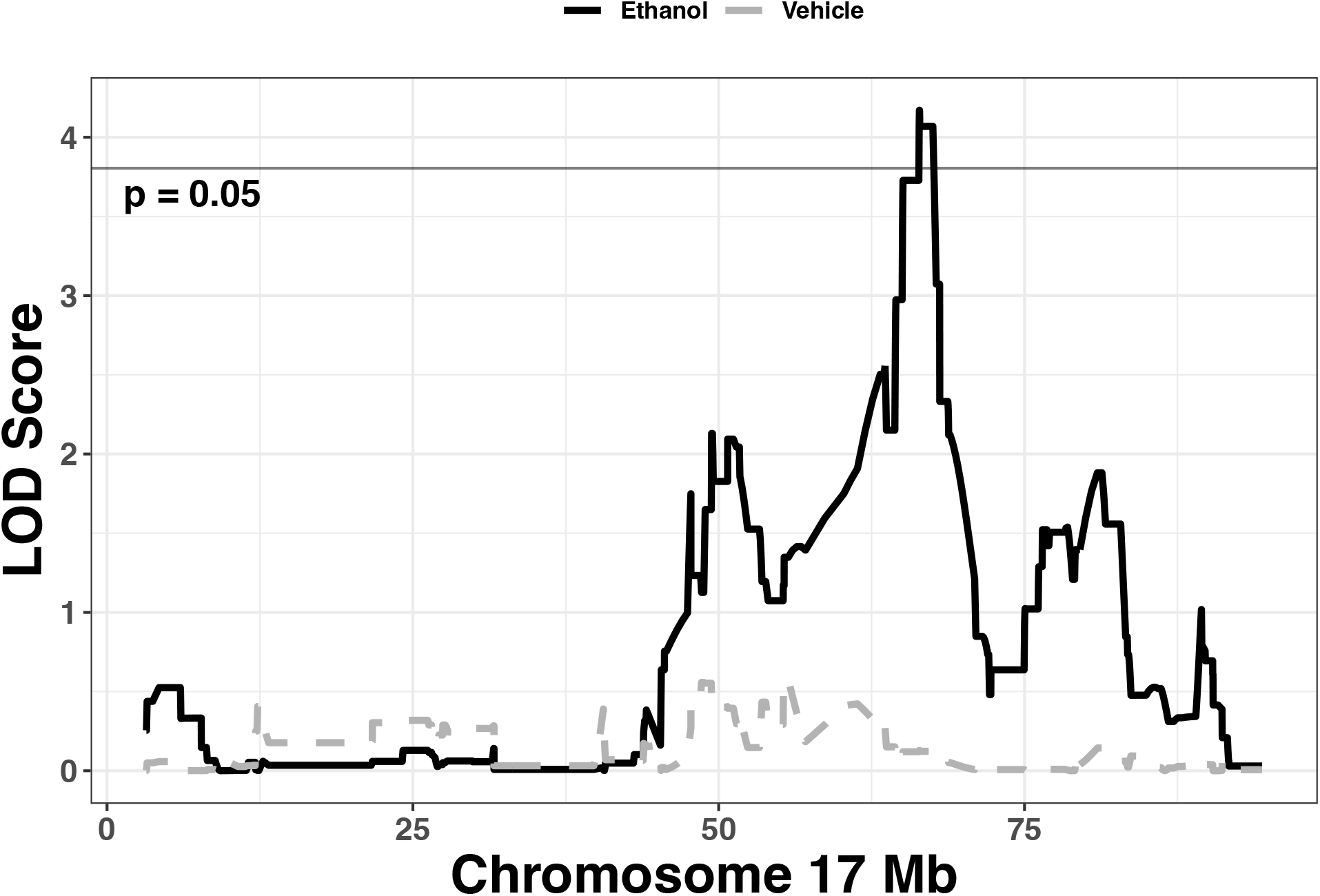
Genome scan of post-ethanol (black) and post-saline (gray) Day 1 hot plate latency across mouse chromosome 17.

**Figure S3.**
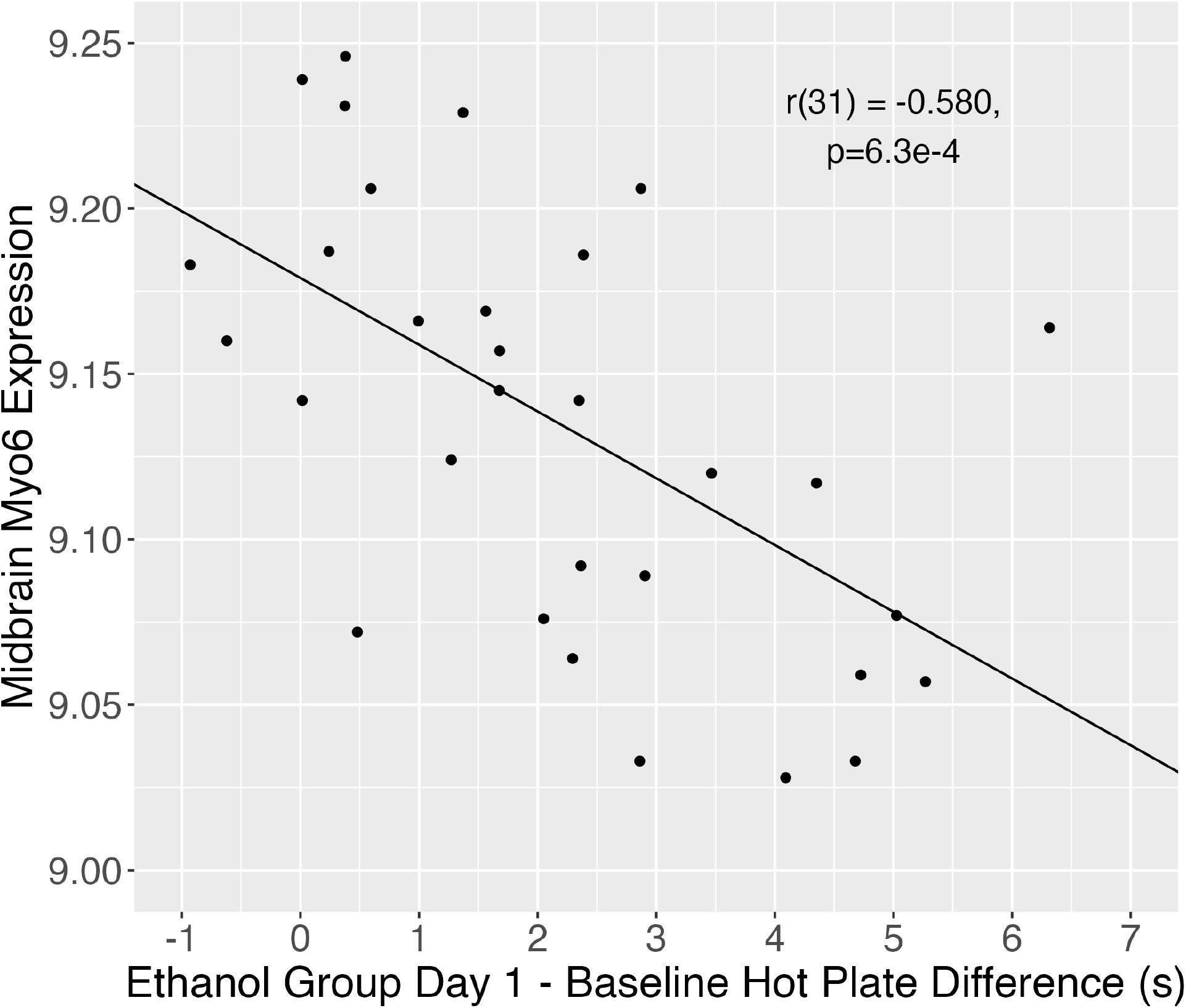
Scatterplot of the correlation between ethanol-specific Day 1 – Baseline hot plate latency difference and midbrain *Myo6* expression.

**Figure S4.**
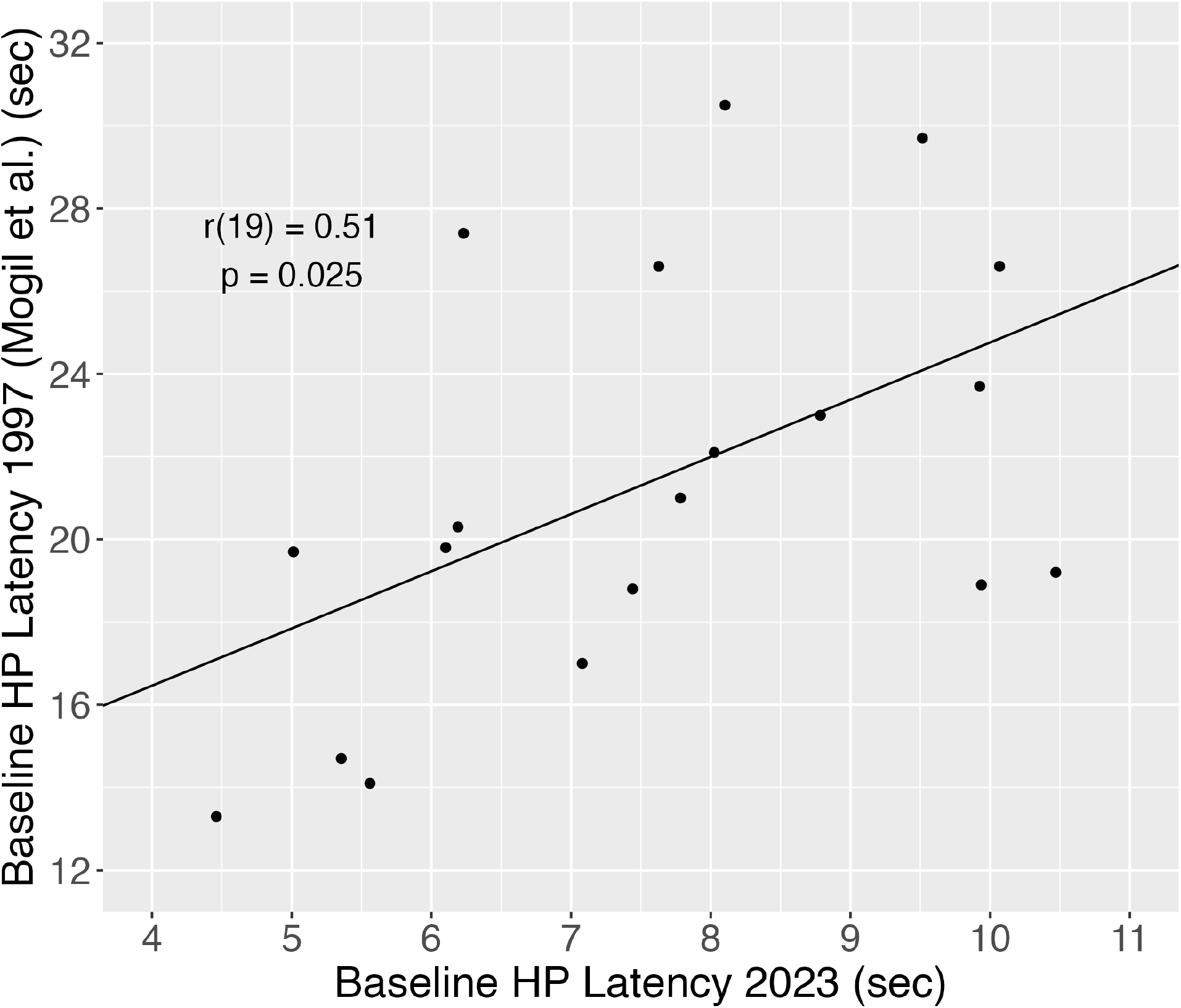
Scatterplot showing significant correlation between the present study basal hot plate latency (x-axis) and baseline latency from a 1997 nociception study by Mogil et al (y-axis).

**Table S1** Protein-coding genes in the 1.5-LOD support intervals of Chromosome 17 (Pehlq1) and Chromosome 9 (Etadq1) bQTLs.

**Table S2** Significant cis-eQTLs for genes in the *Pehlq1* and *Etadq1* support intervals. Transcriptomic databases queried were for prefrontal cortex, midbrain, and nucleus accumbens gene expression.

**Table S3** Spearman rank correlations between prefrontal cortex, midbrain, and nucleus accumbens gene expression levels and Day 1 – Baseline hot plate latency or Day 1 post-ethanol hot plate latency phenotypes.

**Table S4** Top 10,000 Spearman rank correlations between Day 1 post-ethanol hot plate latency or Day 1 – Baseline hot plate latency and publicly available BXD phenotypes on Gene Network.

**Table S5** All behavioral QTL detected across measured ethanol analgesia and hot plate latency phenotypes. Significant, highly suggestive, and suggestive QTL have peak markers with p-values < 0.05, 0.10, and 0.63, respectively. The genomic intervals in the last column were calculated using a 1.5-LOD drop around the peak marker.

**Table S6** Table showing human GWAS results from genomic regions syntenic with mouse bQTL support intervals.

**Table S7.** Heritability estimates for ethanol analgesia and locomotor activity phenotypes in vehicle- and ethanol-receiving mice. MPE=maximum possible effect; LMA = locomotor activity

